# Endosomal trafficking is required for glycosylation and normal maturation of the Alzheimer’s-associated protein sorLA

**DOI:** 10.1101/2020.07.12.199885

**Authors:** Sofie K. Christensen, Yoshiki Narimatsu, Sabrina Simoes, Christoffer K. Goth, Christian B. Vægter, Scott A. Small, Henrik Clausen, Olav M. Andersen

**Affiliations:** Danish Research Institute of Translational Neuroscience (DANDRITE) Nordic-EMBL Partnership, Department of Biomedicine, Høgh-Guldbergs Gade 10, Aarhus University, DK-8000 Aarhus C, Denmark; Copenhagen Center for Glycomics, Department of Cellular and Molecular Medicine and School of Dentistry, University of Copenhagen, Copenhagen N, Denmark; Departments of Neurology and the Taub Institute for Research on Alzheimer’s Disease and the aging Brain, Columbia University, New York, USA

**Keywords:** SorLA, *SORL1*, endosomal trafficking, ectodomain shedding, complex-type *N*-glycosylation

## Abstract

The sorting receptor sorLA encoded by the *SORL1* gene is implicated in Alzheimer’s disease (AD) pathogenesis. Genetic studies have identified AD-associated *SORL1* mutations and the expression of sorLA in AD brains is reported to be reduced. SorLA is a receptor of the retromer trafficking complex and functions at the endosome, and deficiency in sorLA phenocopies the endosomal pathologies found in AD. SorLA undergoes posttranslational modifications and maturation with ultimate ectodomain shedding, however knowledge of these processes remains limited. Here we demonstrate that sorLA exists at the cell membrane in two forms, an immature and a mature form, characterized by distinct *N*-glycosylation profiles. The mature sorLA form has acquired complex type *N*-glycans and is shed from the cell surface by the TACE juxtmembrane cleavage. The immature form of sorLA present at the cell surface is shown to have immature ER-type *N*-glycans (high-mannose type susceptible to endo H) and does not undergo shedding, however, upon endocytosis and recycling to the cell surface via endosomal trafficking pathways the immature sorLA form acquires complex-type *N*-glycans. These results suggest an unusual secretion model for sorLA whereby that immature sorLA first traffics to the cell membrane without acquiring Golgi processing of *N*-glycans, and only upon retrograde trafficking does sorLA acquire normal Golgi maturation of *N*-glycans and become susceptible to TACE regulated shedding. Supportive evidence for this model include a sorLA mutant with deficient endosomal trafficking and *in vivo* studies demonstrating requirement of retromer for sorLA trafficking in the brain of retromer VPS26 deficient mice. Collectively, our study establishes the role endosomal trafficking plays in sorLA’s normal maturation, and point to impaired maturation as a signature of AD-associated sorLA dysfunction.

## INTRODUCTION

*SORL1* was recently established as the fourth autosomal-dominant gene for development of Alzheimer’s disease (AD) (1). A number of genome-wide-association studies (GWAS) have implicated *SORL1* in AD (2–5), and *SORL1* loss-of-function variants are significantly associated with AD in a whole-exome sequencing study (WES) (6). The transmembrane protein encoded by *SORL1* is sorLA, a 250 kDa receptor and its expression in brain is decreased in many AD patients. As a transmembrane protein, sorLA is trafficked via the secretory pathway to the cell surface, from where it internalize by clathrin-coated pits to the early endosome. Once in the endosome, sorLA recycles back to the plasma membrane or undergoes retrograde transport back to Golgi/TGN via the endosomal trafficking complex retromer (7, 8). With particular relevance to AD, studies show that sorLA functions at the endosome (9), where it mediates retromer-dependent trafficking of the Amyloid-Precursor protein (APP) out of the endosome, away for the amyloidogenic cleaving enzymes.

SorLA has 28 predicted *N*-glycosites (7, 10) and some *O*-glycosites (11), and once expressed sorLA is thought to undergo posttranslational maturation as it is transported through trafficking compartments (12, 13). It has previously been observed that sorLA exists as two distinct molecular forms independent of whether expressed exogenously in lysates of cell lines, including NT2 (14), CHO (7), HEK293 (15), SH-SY5Y (16), or expressed endogenously in a leukemia-derived cell lines (17) or in human brain homogenates (18). Previous studies also suggest that it is the mature form of sorLA that is fully functional, and interestingly, the extracellular shedding of sorLA only occurs when sorLA interacts with TACE (TNFα Converting Enzyme) at the cell surface (16, 19). The role of sorLA shedding remains largely elusive, however, the shed ectodomain appears to influence signaling pathways and to regulate cell adhesion (20, 21).

How and where sorLA matures remain unknown, outstanding questions relevant to understanding it’s role in AD pathogenesis. We address these questions by first establishing that both immature and mature sorLA co-exists at the cell surface, where they are characterized by distinct glycosylation profiles. A Golgi-matured glycosylation profile of sorLA is found to be required for TACE cleavage and shedding. While the secretory trafficking pathway is required for delivering immature sorLA to the cell surface, we unexpectedly find that maturation of sorLA only occurs once the immature form is endocytosed and recycled to the cell surface via the endosomal trafficking pathways. We confirm the importance of retromer and endosomal trafficking for sorLA maturation and its relevance to disease, by analysis of a sorLA mutant with impaired endosomal trafficking relevant to AD and by use of a recent retromer deficient mouse model.

## EXPERIMENTAL PROCEDURES

### Cell culture

SH-SY5Y cells were cultured in Dulbecco’s Modified Eagle’s Medium (DMEM – F12, Lonza) containing 10% fetal bovine serum (FBS), 1% penicillin/streptomycin (P/S) and 300 μg/mL zeocin (Invitrogen) at 37°C with 5% CO_2_. Cell lines expressing the mutants of sorLA have previously been described (16, 22, 23). For the production of conditioned medium, cells and serum-free medium were harvested after 48 h. When concentration of medium was needed, this was done using a centrifugal filter with a 50 K cut-off (Amicon Ultra Centrifugal Filter, Merck), at 5,000 rcf in a fixed angle rotor for 10 minutes. CHO cells with knockout (KO) of *Mgat1* (CHO^KOmgat1^) or combinations of KO of *St3gal3/4/6* (CHO^KOst3gal3/4^ and CHO^KOst3gal3/4/6^) were prepared and grown as previously described (24). Phorbol 12-myristate 13-acetate (PMA; Sigma) 100 ng/mL was applied 24 h prior to medium-change left on cells for 0.5, 1, and 2 h before harvesting cells and medium to measure sorLA levels by Western blot analysis. Gm6001 (Sigma) 30 μM was applied for 1 h before medium-change allowing for shedding at indicated time intervals. Dynasore (DYN) and chlorpromazine (CPZ) were applied to cells at a concentration of 50 μM and 30 μM, respectively.

### Animals VPS26b knockouts

All animal procedures and experiments were performed in accordance with national guidelines (National Institutes of Health) and approved by the Institutional Animal Care and Use Committee of Columbia University. Mice were maintained in groups of 5 or less with 12 h on/off light cycles. VPS26b knockout (KO) mice were originally described by Kim *et al*.(25). This line was generated by replacing exons 5 and 6 of the VPS26b gene by a neomycin-resistance cassette. These animals are viable, live into adulthood, and do not display any gross physical abnormalities. Animals from both sex were used in the study. Mice were sacrificed by cervical dislocation, and brain tissue samples were dissected immediately and frozen in dry ice before storage at −80°C. Hippocampi were dissected and analyzed by Western blotting (WB), as described below.

### SDS-PAGE and Western blotting

Loading buffer with 80 mM DTT was added to samples, before heating at 95°C for 5 min and samples were separated by SDS-PAGE using a 4-16% acrylamide gradient gel and blotted onto nitrocellulose membranes (Hybond-C Extra, Amersham Bioscience or Optitran BA-S 85, GE Healthcare). The membranes were incubated in the given primary antibody in a dilution of 1:1000 and appropriate HRP-conjugated secondary antibody (Dako) in a dilution of 1:1500. Recognition of glycosylated sorLA by lectins was carried out applying biotinylated lectins (10 μg/mL; Biotinylated Lectin Kit I and Biotinylated SNA, Vector Laboratories) and peroxidase-conjugated streptavidin (Calbiochem) diluted 1:1500. For analysis SuperSignal^®^ West Femto Maximum Sensitivity Substrate (Thermo Scientific) was used, and blots were developed using LAS-4000 (GE Healthcare). Quantification was performed using densitometry and Multi Gauge v3.2 software. The following antibodies were applied for detection of sorLA (5387, in house rabbit serum), SORCS1 (AF3457; R&D Systems), or calnexin (ab22595, Abcam).

Mouse brain tissue samples were homogenized using a *Glas-Col* homogeneizer in an ice-cold lysis buffer consisting of 20 mM Tris HCl (pH 7.4), 1% Triton X-100, 15 mM NaCl and protease and phosphatase inhibitors (Roche) for protein extraction. After centrifugation at 17,000g for 10 min at 4°C, supernatants were collected and protein concentration was determined using a BCA assay (PIERCE) prior to western blot analysis using the Odyssey blocking buffers (LI-COR Biosciences). Blots were developed using the Odyssey Infrared Imaging System (LI-COR Biosciences) and analyzed using the software program Image Studio Lite Ver 5.0, as specified in the Odyssey software manual. The following antibodies were applied for detection of VPS26b (NBP1-92575, Novus Biologicals) and β-actin (NB600-535, Novus Biologicals).

### Deglycosylation

Endoglycosidase H (endo H; BioLabs), peptide-*N*-glycosidase F (PNGase F) (Roche), neuraminidase (Roche), and *O*-glycosidase (Roche) were used with cell lysates at 50 mU/μg, 100 mU/μg, 0.1 mU/μg, and 50 mU/μg, respectively, and with concentrated conditioned medium (16 μL containing 1% Triton X-100) total amount of enzymes added were 1 U, 2 U, 1 mU, and 1 U endoglycosidase H, respectively. For endo H treatment, denaturation buffer (BioLabs) was added to the lysate, after which G5 buffer (BioLabs) and endo H were added. Sodium-acetate, pH 5.0 in a concentration of 150 mM was added to the samples treated with neuraminidase. The samples with PNGase F and endo H were incubated at 37°C for 1 h. Neuraminidase-treated samples with and without *O*-glycosidase were incubated at RT on.

### Cell surface biotinylation and neuraminidase treatment

SH-SY5Y or HEK293 cells with a confluency of 80% were washed with cold PBS^++^ (PBS with 1 mM MgCl_2_ and 0.1 mM CaCl_2_), and incubated in cold PBS^++^ containing 1 mg/ml membrane impermeable EZ-linked sulfo-NHS-SS-biotin (Thermo Scientific) for 30 min on ice. The reaction was quenched using cold PBS^++^ supplied with 0.1 M glycine (AppliChem). The cells were lysed right away or treated with neuraminidase before lysis. For the neuraminidase treatment each well was supplied with 500 μL of a solution containing 50 mU/mL neuraminidase (Roche Diagnostics) in cold PBS^+++^ pH adjusted to 5.5, and incubated for 1 h at 4°C on ice with gentle agitation. Cells were washed in cold PBS^++^ and lysed using 300 μL RIPA buffer containing protease inhibitor (Complete, Roche). Cell lysates (200 μg) were incubated with 100 μL slurry of Streptavidin beads (GE Healthcare) for 1 h at RT with end-over-end mixing. Beads were washed five times with RIPA buffer before protein was eluted with loading buffer containing 60 mM DTT at 95°C for 5 min. The total protein fraction and the cell surface fraction were analyzed with SDS-PAGE and Western blotting.

### Immunocytochemistry

Cells were seeded on poly-L-lysine coated coverslips and after cultivation fixed in 4% paraformaldehyde (Sigma) 15 min. Cells were permeabilized n PBS with 0.1% Triton X-100 (AppliChem) and incubated in 10% FBS for 30 min. The cells were then incubated overnight at 4°C in primary antibody dissolved in a 10% FBS solution. Sol-sorLA antibody (catalog #5387, Dako) was used in a concentration of 1:500. The cells were washed in PBS with or without 0.1% Triton X-100 and incubated in secondary antibody Alexa flour 488 donkey anti-rabbit (Invitrogen) in a concentration of 1:500 in 1 h at RT in a dark box. Following this the cells were washed in ddH2O containing Hoechst nucleus staining in a concentration of 1:10,000. The coverslips were mounted using DAKO Flouromount (Dako) and visualized with Zeiss confocal LSM 780 with C-Apochromat 63x/1.20 W Korr M27 objective. The software used for image processing was ZEN Black (Zeiss).

### Pulse-chase experiments

SH-SY5Y cells expressing sorLA^WT^, sorLA^KKLN^, or sorLA^ΔCD^ with a confluency of about 90% were washed twice in methionine, cysteine and glutamine free media (Sigma) supplied with 1% Glutamax (Sigma) and 1% P/S. The cells were incubated for 30 min at 37°C in 1.5 mL media added 2% dialyzed FBS with or without 30 μM CPZ, or 1 μg/mL Tun. After 30 min this solution was removed and the cells were metabolically labeled for 40 min at 37°C in 1.5 mL medium with 1% P/S, 2% dialyzed FBS and Pro-mix containing L-[^35^S]cysteine and L-[^35^S]methionine (Amersham Bioscience) with or without CPZ or Tun. After the pulse, the medium with Pro-mix was removed and the cells were washed and incubated in the same medium with or without CPZ or Tun. Cells and medium were harvested at the given chase times. The medium was centrifuged and protease inhibitor was added. To immunoprecipitate sorLA proteins 70 μL GammaBind Beads (GE Healthcare) coated with sol-sorLA antibody was used for each sample. 200 μL of the cell lysates or 1.5 mL of the medium were added to the coated beads and incubated over night with end-over-end mixing. The beads were washed in 0.05% Tween and the proteins eluted in 35 μL sample buffer with DTT and heated to 95°C for 5 min. Following SDS-PAGE analysis the gels were fixed in fixation buffer for 30 min and incubated in DMSO (Merck) for 2 × 30 min. The gels were incubated in 22% 2,5-diphenyloxazole (PPO) in DMSO for 30 min and the gel was dried overnight. The gel was analyzed with radio-fluorography using FLA-3000 (Fujifilm).

### In vitro TACE assay

A peptide spanning the stem region of sorLA (amino acids 2,103 to 2,137) was designed and purchased (NeoBiolab Inc). *In vitro* cleavage was assayed by adding 600 nM recombinant TACE (Enzo Life Sciences), using 10 μg of peptide substrate in a total volume of 25 μl. Reactions were performed in 25 mM Tris-HCl, pH 9 and incubated at 37°C. Product development was evaluated by MALDI-TOF-MS and cleavage sites were determined by comparing the mass of the cleavage fragments with the theoretical masses.

### Statistical analysis

All statistical tests were done using the program Prism GraphPad version 6.0. For comparison between two different datasets a two-tailed unpaired Student’s *t*-test or non-parametric Mann-Whitney *t*-test were used. A *p* value above 0.05 was not considered significant (ns), whereas *p* < 0.05, (*),*p* < 0.01 (**), and *p* < 0.001 (***) were considered as significant different. Error bars represent standard errors of the mean.

## RESULTS

### Characterization of the mature and immature forms of sorLA

We previously observed that sorLA proteins from total cell homogenates migrate as two distinct bands by SDS-PAGE and WB analysis, while sorLA from culture medium migrates as a single band with a mobility in between these forms (7, 14–17). This was true for most cell types including SH-SY5Y, and for endogenous sorLA as well as with exogenous expressed constructs (Fig. 1A,B). To explore the molecular basis of this observation, we used cell surface biotinylation to assess the molecular forms present at the cell surface of SH-SY5Y as well as HEK293 cells using expression of full coding sorLA^WT^ constructs (Fig. 1C,D). Surprisingly, we could clearly identify both the upper and the lower variant in the pool of cell surface receptors. As a control the blots were incubated with an antibody that detects the ER protein calnexin, which was not present among the biotinylated proteins (Fig. 1C). SorLA has 28 predicted *N*-glycan sites and changes in glycosylation present obvious explanations for multiple migrating bands (7, 10, 14, 23). We also recently identified *O*-glycosites on sorLA (12, 13). Neuraminidase treatment of surface biotin-labelled proteins bound on Streptavidin-conjugated beads prior to SDS-PAGE and WB analysis, revealed that the two sorLA bands collapsed into one band migrating as the lower form found in lysates using both SH-SY5Y and HEK293 cells (Fig. 1D).

**Fig. 1.**
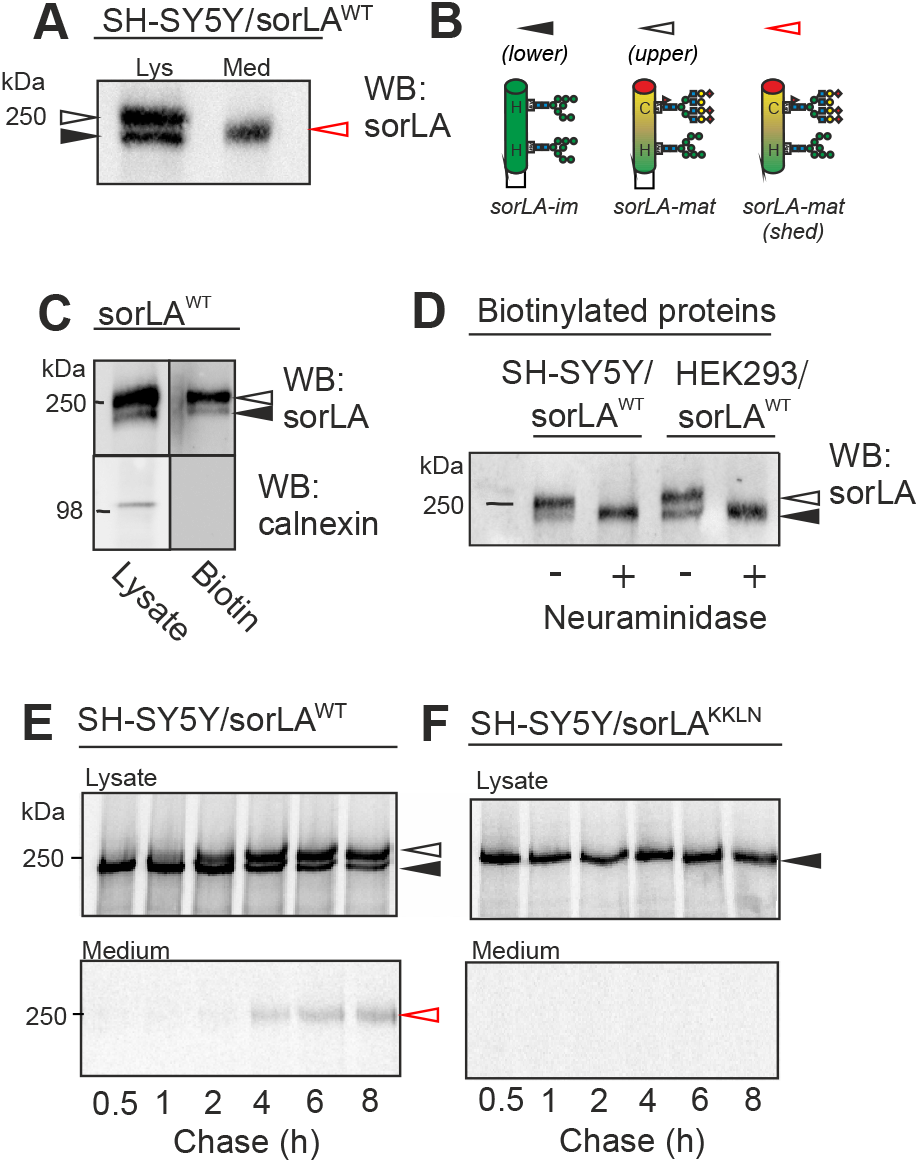
Two sorLA variants found at the cell surface but only one is shed. **(A)** WB analysis of lysate and conditioned medium from SH-SY5Y cells stably transfected with sorLA^WT^. **(B)** Schematic representation of proposed *N*-glycosylation variants for two molecular species of sorLA (mature: arrow open, and immature: arrow filled) in lysates and one in medium (mature shed: arrow red). The high molecular weight form (upper band; sorLA-mat) carries a mixture of mature complex-type (C) and high-mannose (H) *N*-glycans, whereas the low molecular weight form (lower band; sorLA-im) only carries immature high-mannose *N*-glycans. **(C)** Cell surface proteins from SH-SY5Y cells expressing exogenous sorLA^WT^ were labeled with biotin and precipitated using Streptavidin coated beads. The biotinylated proteins together with total lysates were analyzed using SDS-PAGE and WB for sorLA and calnexin (ER marker). **(D)** Eluates of biotinylated cell surface proteins from SH-SY5Y and HEK293 cells expressing exogenous sorLA^WT^ treated (+) or not (-) with neuraminidase after precipitation using Streptavidin coated beads as indicated. **(E, F)** SH-SY5Y cells expressing either sorLA^WT^ or sorLA^KKLN^ were subjected to a [^35^S] pulse-chase protocol in order to follow the maturation over time. Cells were harvested at the indicated time points, and sorLA protein from lysate and medium was immunoprecipitated, separated by SDS-PAGE analysis, and visualized by radio-fluorography. Signals in WB for lysates for sorLA-mat and sorLA-im are indicated with white and black arrowheads, and for shed sorLA in medium with a red arrowhead, respectively.

We next used metabolic [^35^S]Cys-Met pulse-chase labeling in SH-SY5Y cells and followed sorLA^WT^ by immunoprecipitation and SDS-PAGE autoradiography (Fig. 1E). In the total cell lysates, the lower band of sorLA^WT^ was visible from the beginning (0.5 h) and decreased in intensity throughout the chase period, while the upper band appeared at 2-4 h and increased in intensity up to 6 h. In striking contrast, in the culture medium only a single band was detectable, and the appearance of this band coincided with the timing of the upper band in lysates. We included a sorLA variant carrying a fused C-terminal KKLN tetrapeptide (sorLA^KKLN^) leading to ER-retention as a control (23), and this demonstrated that the ER-retained form did not undergo maturation to the upper band in total cell lysates, neither did it undergo shedding into the medium (Fig. 1F). Furthermore, direct comparison of the migration of bands with sorLA^WT^ and sorLA^KKLN^ by WB analysis, confirmed that the single band for sorLA^KKLN^ migrated similar to the low immature band of sorLA^WT^ (*Supporting Information*, Fig. S1). Combined, these results suggested that the upper band was representative of a mature *N*-glycoforms of sorLA with complex-type *N*-glycans, while the lower band was representative of an immature *N*-glycoform likely representing high-Man without complex-type sialylated glycans, as illustrated in Figure 1B. This hypothesis would suggest that while both *N*-glycoforms of sorLA reaches the cell surface, only the glycoform with complex-type *N*-glycans undergoes shedding.

To test this we performed deglycosylation with Endo H and PNGase F of cell lysates and media from SH-SY5Y cells expressing sorLA^WT^ (Fig. 2A,B). PNGase F treatment collapsed the two sorLA^WT^ bands in cell lysates into a single band migrating 20-30 kDa faster confirming that both bands represent sorLA with multiple *N*-glycans (Fig. 2A). Interestingly, both of these forms were also sensitive to treatment with Endo H, however, the upper sorLA band was only partially sensitive since the product did not co-migrate with the product of PNGase F. The lower band in contrast shifted migration to a similar position as after PNGase F treatment. Treatment with a broad neuraminidase confirmed that only the upper band was sensitive and the treatment collapsed the two bands into one migrating as the lower band. Combining neuraminidase and *O*-glycanase did not further shift migration, suggesting that *O*-glycans do not substantially affect SDS-PAGE mobility of sorLA in SH-SY5Y cells (Fig. 2A). The same analyses with the culture medium of SH-SY5Y cells expressing sorLA^WT^ confirmed that the shed upper band was only partially sensitive to endo H but sensitive to neuraminidase treatment (Fig. 2B). This finding was in agreement with the presence of sialic acids only on the upper and shed band as shown by recognition by the lectin SNA (*Supporting Information*, Fig. S2).

**Fig. 2.**
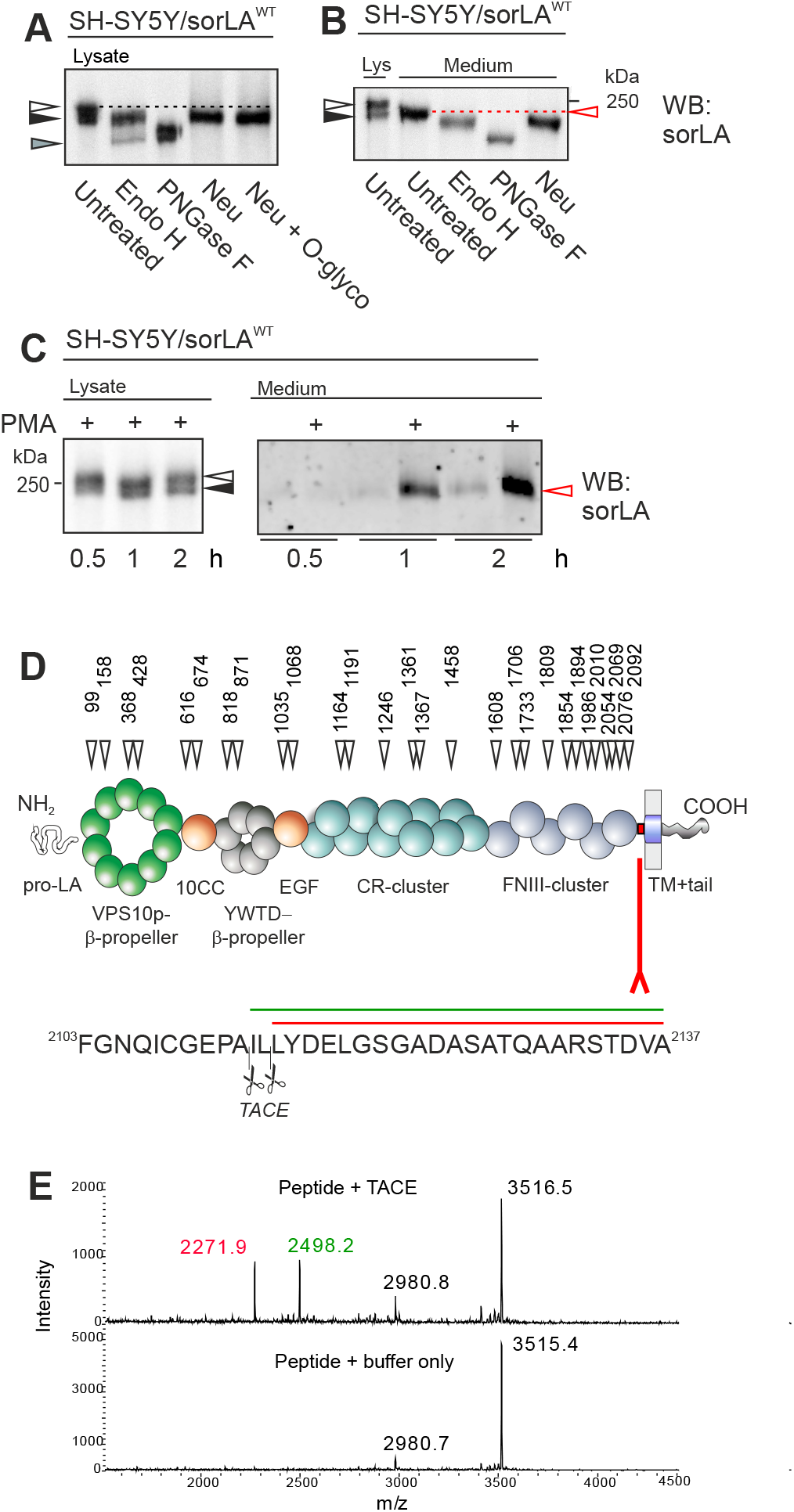
Two distinct sorLA cell surface variants differ by the presence of complex-type *N*-glycans. **(A, B)** Lysates and medium from SH-SY5Y cells expressing exogenous sorLA^WT^ were enzymatically digested with Endo H, PNGase F, and either neuraminidase (Neu) alone or in combination with *O*-glycosidase. Protein samples were subsequently separated by SDS-PAGE and analyzed by WB for sorLA. Both the lysate form of sorLA-mat (A) and the shed form of sorLA (B) migrate faster after Neu exoglycosidase treatment. Signals for sorLA-mat (upper band) and sorLA-im (lower band) in lysates are indicated with white and black arrowheads, respectively, while shed sorLA represented by a red arrow. The grey arrowhead indicates migration of the nascent sorLA protein in lysate with no *N*-glycans. **(C)** SH-SY5Y cells expressing exogenous sorLA^WT^ were treated with phorbol 12-myristate 13-acetate (PMA) for the indicated time periods before medium and cells were harvested, followed by SDS-PAGE and WB testing for effects on sorLA shedding. **(D)** Schematic domain representation of full-coding sorLA. Triangles indicate the position of *N*-X-S/T consensus motifs for *N*-glycosylation (28 sites) and numbers correspond to the Asn position in the human sorLA protein sequence. The juxta-membrane stem region (aa 2,103-2,137) is indicated by a red fork. **(E)** MALDI-TOF analysis of *in vitro* TACE assay with a sorLA stem peptide (mass 3,516 Da) after 6 h reaction time. The observed fragments were matched to cleavage sites as indicated by scissors and colored bars in the primary sequence.

It was previously shown that TACE is the enzyme responsible for sorLA shedding (14, 26, 27), however, the regulation of this process is so far incompletely understood (28, 29). We therefore tested the effect of the TACE activator PMA (30–32) on sorLA^WT^ shedding in SH-SY5Y cells (Fig. 2C). There was no effect of PMA treatment on the expression levels or forms of sorLA in total cell lysates, however, there was a strong induction of shed receptor fragment in the medium as early as 1 h after PMA treatment. In parallel experiments we also tested ionomycin, which specifically activates the related sheddase ADAM10 (31). However, we did not see any significant effect for sorLA shedding upon treatment of SH-SY5Y cells with ionomycin (not shown). These data confirm that TACE is the major enzyme responsible for sorLA shedding also in SH-SY5Y cells, and demonstrated again that only the single mature sorLA glycoform with complex-type *N*-glycans is shed.

To explore the nature of the cleavage site for TACE on sorLA we used an *in vitro* cleavage assay with a peptide derived from the juxtamembrane region of sorLA (amino acids 2103-2137) monitored by MALDI-TOF analysis (Fig. 2D). TACE cleaved at two distinct positions in the stem region (^2111^PA|IL|LY^2116^) (Fig. 2E). The immediate adjacent region does not appear to contain obvious posttranslational modifications including sites for *N*- or *O*-glycosylation, although we note that the AILL motif is adjacent to a Tyr site in YDE sequon that may undergo e.g. sulfation or phosphorylation (Fig. 2D).

We next tested if *N*-glycosylation of sorLA (schematic illustration in Fig. 3A) was required for transport through the secretory pathway to the cell surface using tunicamycin treatment of SH-SY5Y cells transfected with sorLA^WT^. Tunicamycin inhibits the biosynthesis of the precursor oligosaccharide substrate of the oligosaccharyltransferase, and hence blocks addition of *N*-glycans to nascent polypeptides. WB analysis of cell lysates confirmed that tunicamycin inhibited addition of *N*-glycans to sorLA^WT^ (Fig. 3B). Cell surface biotinylation experiments performed in the presence of tunicamycin further showed that unglycosylated sorLA^WT^ reached the cell surface (Fig. 3B), indicating that sorLA^WT^ traffics efficiently to the plasma membrane in the absence of *N*-glycans. We continued with [^35^S] metabolic labeling of SH-SY5Y cells expressing sorLA^WT^ with and without tunicamycin (Fig. 3C). The pulse-chase study demonstrated that tunicamycin totally abolished cleavage and shedding of sorLA^WT^.

**Fig. 3.**
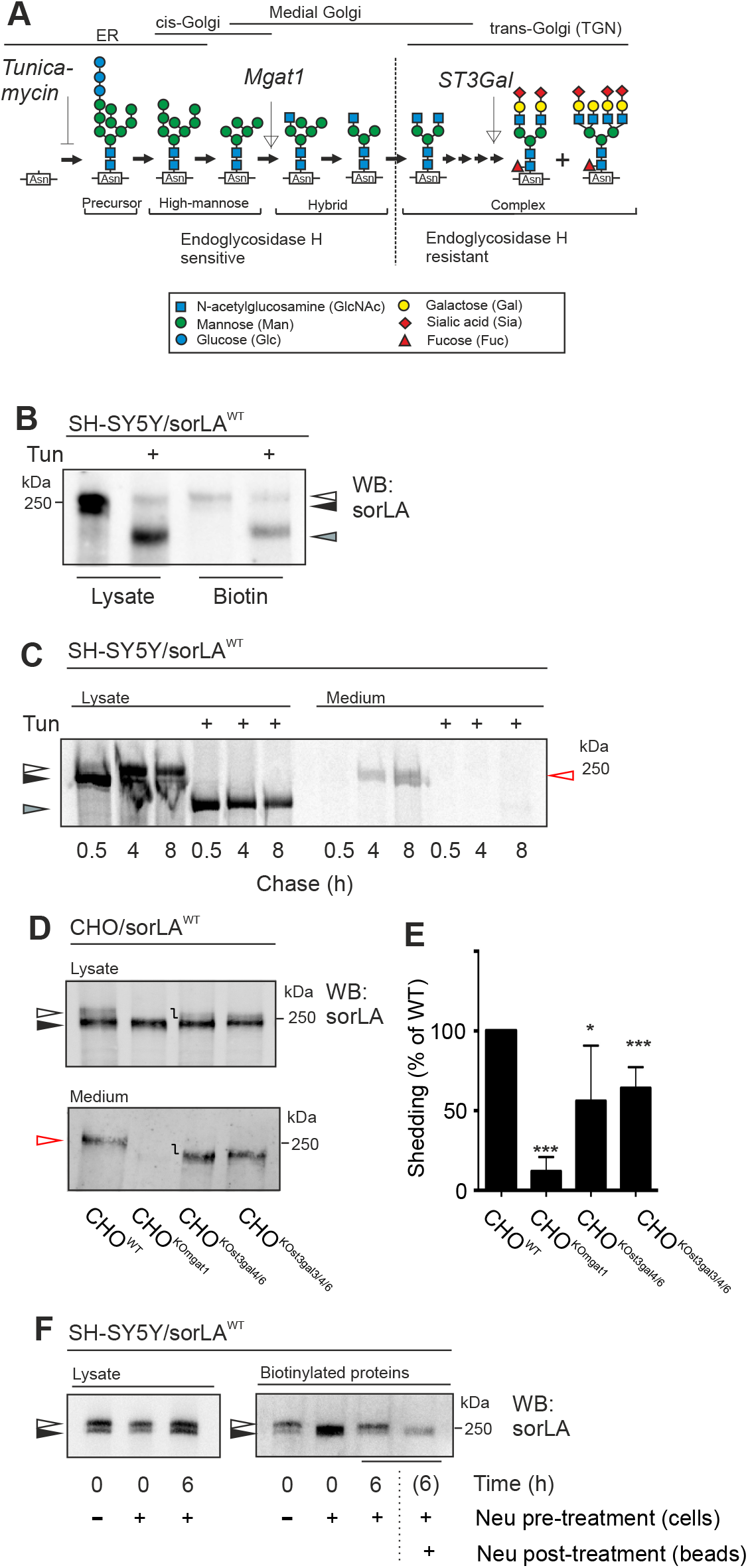
SorLA needs complex-type *N*-glycosylation for shedding. **(A)** Schematic representation of classical *N*-glycan maturation in the secretory pathway. Tunicamycin blocks initiation of protein *N*-glycosylation, whereas MGAT1 is required for conversion of high-mannose glycans and ST3Gal sialyltransferases for capping of the glycans as indicated. Designations for monosaccharides follow the Consortium for Functional Glycomics. **(B)** SH-SY5Y cells expressing exogenous sorLA^WT^ were treated with tunicamycin (1 μg/mL) for 24 h prior to biotinylation of cell surface proteins. The biotinylated proteins together with total lysates were analyzed using SDS-PAGE and WB for sorLA. **C)** SH-SY5Y cells expressing exogenous sorLA^WT^ were subjected to a 40 min [^35^S]-labeling and chased for the indicated time points in absence or presence (+) of tunicamycin. SorLA proteins from cell lysates and medium were immunoprecipitated, and subsequent analyzed by SDS-PAGE and radio-fluorography. **(D)** Wild type CHO (CHO^WT^) or CHO cells with targeted deletion of *Mgat1* (CHO^KOmgat1^), *St3gal4* and *St3gal6* (CHO^KOst3gal4/6^), or *St3gal3, St3gal4*, and *St3gal6* (CHO^KOst3gal3/4/6^) were transfected with sorLA^WT^ and shedding efficiency determined by WB analysis of cell lysates and conditioned medium. **(E)** Quantification of shed sorLA from three independent experiments as illustrated in panel D. Signals for sorLA-mat (upper band) and sorLA-im (lower band) in lysates are indicated with white and black arrowheads, respectively, while shed sorLA represented by a red arrow. The grey arrowhead indicates migration of the nascent sorLA protein in lysate with no *N*-glycans. **(F)** SH-SY5Y cells expressing exogenous sorLA^WT^ were subjected to surface biotinylation (time 0 h) and treatment with neuraminidase (Neu) to remove sialic acids on surface located sorLA. Following cultivation at 37°C for 6 h total cell lysates were subjected to precipitation on Streptavidin beads (time 6 h) and analyzed by WB for detection of sorLA species. Note that surface biotinylated sorLA at time 0 h after neuraminidase treatment co-migrates with the lower migrating sorLA-im, and at time 6 h regains co-migration with the sorLA-mat upper band. To demonstrate that the surface biotinylated sorLA species detectable after 6 h indeed had regained sialic acids the streptavidin precipitate was subjected to neuraminidase again (Neu post-treatment, beads), and in agreement with this a clear shift in mobility was observed.

To further dissect the need of complex-type *N*-glycans for sorLA shedding, we used glycoengineered CHO cells with knockout (KO) of the *mgat1* gene that encodes the key enzyme initiating complex-type *N*-glycosylation (see Fig. 3A) (33). Expressing sorLA in CHO^WT^ recapitulated that sorLA migrates as two bands with only the upper band shed into the medium, while sorLA expressed in CHO^KOmgat1^ without complex-type *N*-glycans migrated as the lower single band (Fig. 3D). More importantly, the CHO^KOmgat1^ cells failed to shed sorLA into the medium in agreement with our previous studies. We then proceeded to test the role of sialic acid capping of complex-type *N*-glycans by using CHO cells with selective KO of the predominant α2-3sialyltransferases, ST3GAL3, 4, and 6, involved in capping of *N*-glycans but not *O-* glycans (33). Expressing sorLA in CHO^KOst3gal3/4^ and CHO^KOst3gal3/4/6^ produced only lower migrating sorLA bands confirming that the upper band of sorLA represents the glycoform with complex-type *N*-glycans capped by sialic acid residues (Fig. 3D), in agreement with our studies showing sensitivity to neuraminidase treatment (Fig. 1D). However, interestingly, the KO of sialylation capacity for *N*-glycans did not substantially affect the shedding of sorLA (Fig. 3DE). This indicates that while complex-type *N*-glycans are required for shedding of sorLA, the inherent capping step of *N*-glycans by sialic acids does not appear to be required for shedding.

Collectively, these studies revealed that glycosylation of sorLA is not a required step for delivery to the cell surface, while complex-type *N*-glycans, but not necessarily sialylation, is required for TACE-mediated ectodomain shedding.

### SorLA maturation occurs in endosomal trafficking pathways

*N*-glycans undergo trimming of high-Man structures in the ER before the MGAT1 enzyme initiates complex-type structures in cis-Golgi (Fig. 3A), and these are then sequentially branched, elongated and capped by sialic acids throughout the secretory pathway (34). However, a number of studies have shown that cell surface proteins during recycling to endosome or late Golgi compartments/TGN may undergo further glycosylation, and especially well documented is resialylation. Among the best characterized examples are the two mannose-6-phosphate receptors that during recycling to Golgi compartments are sialylated (35–38). However, also the low-density lipoprotein receptor (LDLR) (37), transferrin receptors (TFRC) (39, 40), as well as Episialin (41) and the serine peptidase dipeptidylpeptidase IV (42), are known to undergo retrograde sorting to Golgi/TGN compartments and become resialylated. While we found that sorLA requires complex-type *N*-glycans, but not necessarily sialylation, for TACE-dependent ectodomain shedding (Fig. 3DE), we wanted to explore if sorLA after desialylation at the cell surface can undergo resialylation during recycling (Fig. 3F).

We utilized a well-established assay previously used to demonstrate that cell surface receptors undergo resialylation upon recycling to the late Golgi/TGN (35, 42, 43). SH-SY5Y cells expressing exogenous sorLA^WT^ were cell surface labelled at 0°C with a membrane impermeable biotinylation reagent and sialic acids removed by treatment with neuraminidase (time 0 h), followed by cultivation for 6 h at 37°C. Analysis of total cell lysates by SDS-PAGE WB analysis showed that cell surface biotin-labelled sorLA migrated as the two characteristic bands before and after the 6 h culturing regardless of whether sialic acids were removed before culturing (Fig. 3F). We further confirmed that the surface-biotinylated sorLA after 6 h culturing contained sialic acids by retreating the Streptavidin beads with neuraminidase, which resulted in the expected collapse of upper and lower bands into a single migrating band (Fig. 3F). Note that the major pool of sorLA (>90%) is located intracellularly (7), and thus treatment with neuraminidase had no apparent effect on migration of sorLA in total lysates (Fig. 3F).

At the cell surface sorLA undergoes clathrin-mediated endocytosis (7, 8). To explore the role of internalization and retrograde trafficking for acquisition of complex-type *N*-glycosylation, we tested sorLA maturation in cells treated with pharmacological endocytosis inhibitors. We choose Dynasore (DYN) and Chlorpromazine (CPZ) to block dynamin-dependent endocytosis and delivery from coated vesicles to endosomes, respectively (44) (Fig. 4A). In untreated cells, sorLA^WT^ was mainly located perinuclear in untreated SH-SY5Y cells, in agreement with previous studies showing predominant Golgi/TGN localization (7, 22). DYN treated cells showed sorLA^WT^ expression at the cell surface, and CPZ treatment resulted in accumulation of sorLA^WT^ in endososomal-like vesicles (Fig. 4B). Quantification of the two sorLA forms by WB of total cell lysates after surface biotinylation of cells with and without drug treatment showed that the majority (65 ± 2 %) of sorLA^WT^ existed as the mature variant in non-treated cells (Fig. 4C). In contrast, when endocytosis and retrograde trafficking were blocked by either DYN or CPZ treatment, significant (p=0.0039 and p=0.0018) decrease in receptor maturation were found, with only 49 ± 3 % or 40 ± 4 % of the cellular receptor pool converted to the mature variant (Fig. 4D). As expected, DYN increased receptor expression at the cell surface by 2.6-fold (p=0.050), whereas CPZ did not significantly change the overall level of sorLA^WT^ located at cell surface as determined by quantification of WB analysis in three independent biotinylation experiments (187 ± 46 % relative to 100 % for non-treated cells) (Fig. 4D). This finding is in agreement with CPZ specifically blocking trafficking away from endosomes and not affecting the initial step of internalization from the plasma membrane. We also observed a consistent change in the ratio between the upper and the lower migrating bands of surface biotinylated sorLA^WT^ for both conditions, indicating that both DYN and CPZ treatment inhibit maturation of sorLA *N*-glycans from high-mannose to the complex-type. The use of two different pharmacological inhibitors, both leading to a similar result, clearly support the conclusion that sorLA maturation depends on endocytosis, and decreases the possibility that our findings are affected by unspecific, off-target effects.

**Fig. 4.**
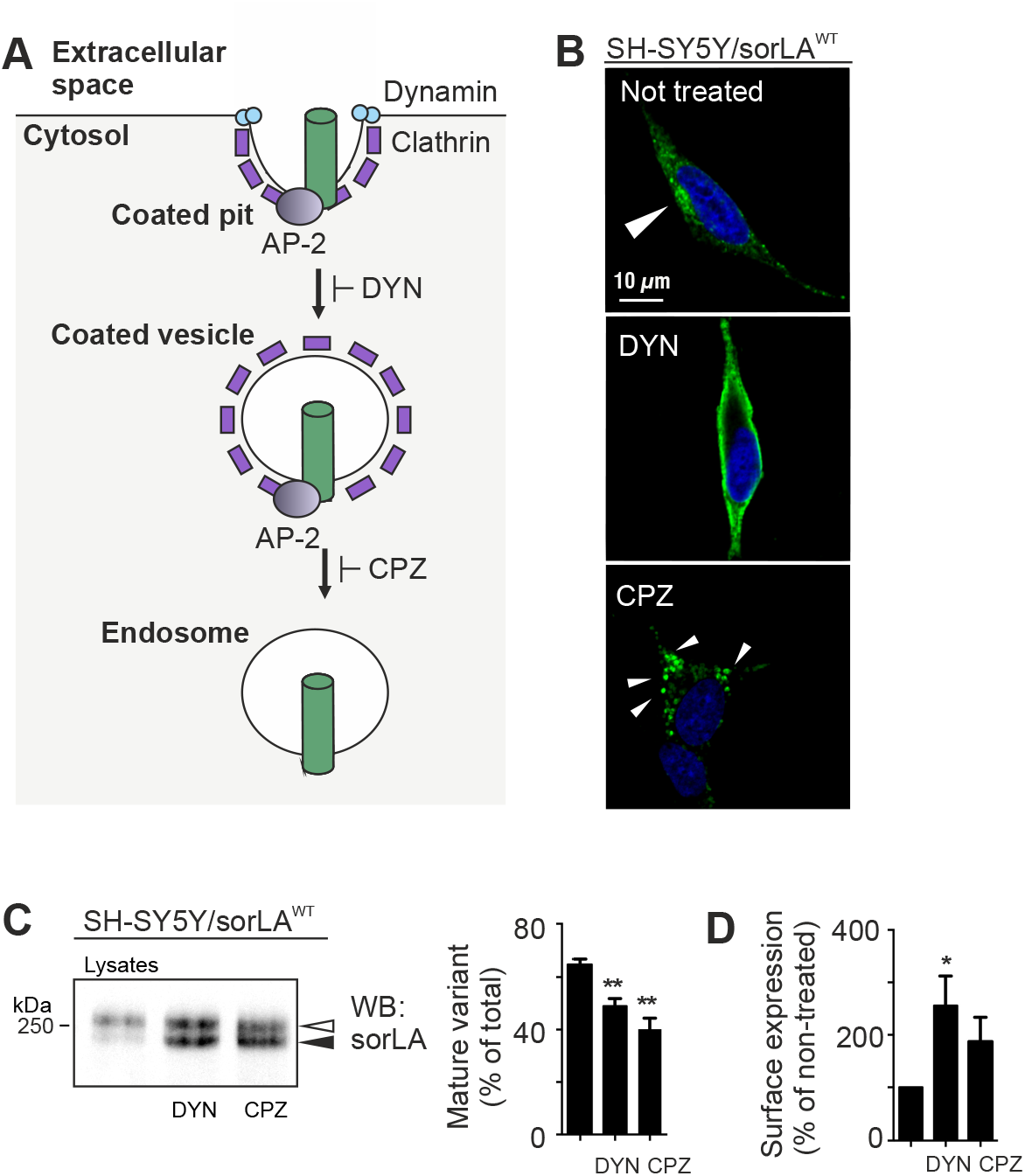
DYN and CPZ inhibit sorLA^WT^ complex-type *N*-glycan formation in the endocytic pathway. **(A)** Schematic representation of the action of dynasore (DYN) and chlorpromazine (CPZ) that block dynamin-dependent endocytosis and the conversion from coated vesicles to endosomes during clathrin-coated internalization, respectively. **(B)** Confocal microscopy displaying the localization of sorLA^WT^ in SH-SY5Y cells treated with DYN (center) or CPZ (bottom) for 24 h or non-treated cells (top). The large white arrowhead indicates perinuclear localization of sorLA^WT^ in non-treated cells. The small white arrowheads indicate punctuate (i.e. endosome-like vesicular) localization of sorLA^WT^ in cells treated with CPZ (scale bar = 10 μm). **(C)** SH-SY5Y cell expressing exogenous sorLA^WT^ were treated with DYN or CPZ for 24 h prior to biotinylation of surface proteins. Total lysates and biotinylated proteins were analyzed using SDS-PAGE and Western blotting for sorLA. The signals for the mature and the immature sorLA^WT^ in samples from total lysates were determined by densitometric scanning of Western blots from three independent experiments and.the level of maturation (expressed as the level of the upper variant relative to the sum of both variants) is shown in bar graph, as mean ± S.E.M. **(D)** Bar graph representing the amount of sorLA^WT^ on the cell surface of both variants relative to the total level of sorLA on the plasmamembrane on cells with no treatment.

We next followed the maturation of sorLA^WT^ using [^35^S]-metabolic labeling of newly synthesized receptors in SH-SY5Y cells expressing sorLA^WT^ with and without CPZ treatment (Fig. 5A, B). Without added CPZ the mature variant was formed ~2-4 h after initiation of the chase period (Fig. 5A), as shown above (Fig. 1E). Densitometric quantification of gels from three independent experiments showed that after 8 h chase 58% of sorLA^WT^ was present as its mature variant, and only 28% was left as the immature high-Man form. Interestingly, this time course is similar to those reported for recycling and sialylation of TFRC, LDLR, and IGF2R (35–37, 39), suggesting that sorLA follows a similar sorting pathway. In contrast, CPZ treatment strongly compromised conversion from the lower to the upper band, and the majority of sorLA remained as the immature variant even after 8 h chase (Fig. 5B), which supports the hypothesis that retrograde transport is required for complex-type *N*-glycosylation of sorLA.

**Fig. 5.**
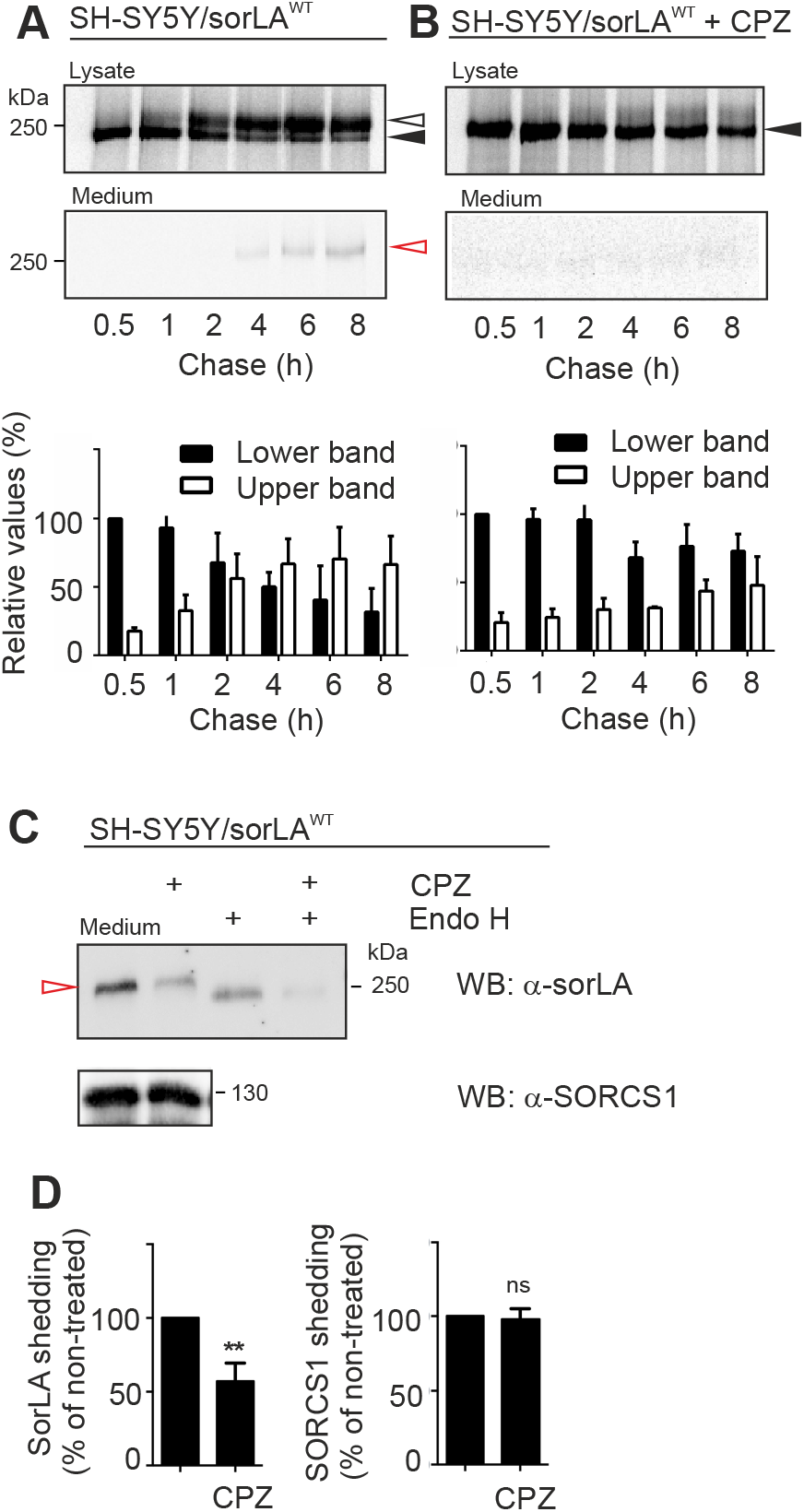
CPZ decreases sorLA shedding. **(A, B)** SH-SY5Y cells expressing exogenous sorLA^WT^ were labelled with [^35^S] for 40 min and chased up to 8 h in the absence (A) or presence (B) of 30 μM CPZ. The cells were then harvested at the indicated time points, before sorLA proteins were immunoprecipitated and subsequent analyzed by SDS-PAGE and radio-fluorography. A red arrowhead indicates migration of shed sorLA in the medium. The signals for the two cellular sorLA forms corresponding to mature (white) and immature (black) were quantified using densitometry of gels from three independent experiments, and plotted as relative to the signal for immature sorLA at 0.5 h (set to 100%). **(C)** The amount of shed ectodomain at steady-state conditions in the absence or presence (+) of CPZ was determined by WB for sorLA^WT^ and the non-affected SORCS1 in 48 h conditioned medium. Treatment of medium with Endo H confirmed that shed sorLA carried high-mannose *N*-glycans. **(D)** Quatifications of shed sorLA and SORCS1 and the bar graph represents values from four independent experiments, with values for the mean (± S.E.M).

A similar result was obtained when looking at non-labeled cells grown for 24 h (Fig. 5C), where quantification of WBs from three independent experiments confirmed a significant 43 ± 12 % reduction (p=0.013) in the shedding of sorLA^WT^ in the presence of CPZ relative to non-treated cells. Endo H digestion confirmed that shed sorLA was immature carrying high-man *N*-glycans under these conditions (Fig. 5C, D). As a control, CPZ had no effect on shedding of SORCS1, another VPS10p-domain containing receptor related to sorLA expressed in SH-SY5Y cells (Fig. 5D). From these experiments, we conclude that sorLA endocytosis and endosomal recycling is required for the full maturation of sorLA and its shedding.

### Modelling Alzheimer’s-associated mutations impairs its maturation

We next tested maturation of sorLA mutants with introduced sorting defects by expression in CHO and SH-SY5Y cells (described in *Supporting Information*, Fig. S3). SorLA^ΔCD^ is truncated by the most C-terminal 46 amino acids in the cytoplasmic domain (CD), and previous studies have found that the CD deleted receptor traffics to the cell surface from where it cannot internalize due to the lack of endocytosis signals in its CD (22, 23, 45). SorLA^ΔCD^ also displayed as a doublet band in WB analysis (Fig. 6A), but with a more intense lower band and strong ectodomain shedding of the CD tail-less receptor as previously noticed (22, 23, 45), suggesting that the lack of the upper band in lysates is a direct consequence of the enhanced shedding. This was confirmed by inhibition of TACE activity using Gm6001 (Fig. 6B), as well as pulse-chase assays (Fig. 6C) and WB analysis (Fig. 6D). It is noteworthy that shedding of sorLA^ΔCD^ appeared to occur much faster and efficient than sorLA^WT^, suggesting that the tail-less variant follows a different and more direct route to the cell surface without the need for internalization before being shed. We propose that this suggests that the cytosolic tail of sorLA mediates interactions with cargo proteins during its first pass to the cell surface that may prevent the normal Golgi maturation to complex-type *N*-glycans.

**Fig. 6.**
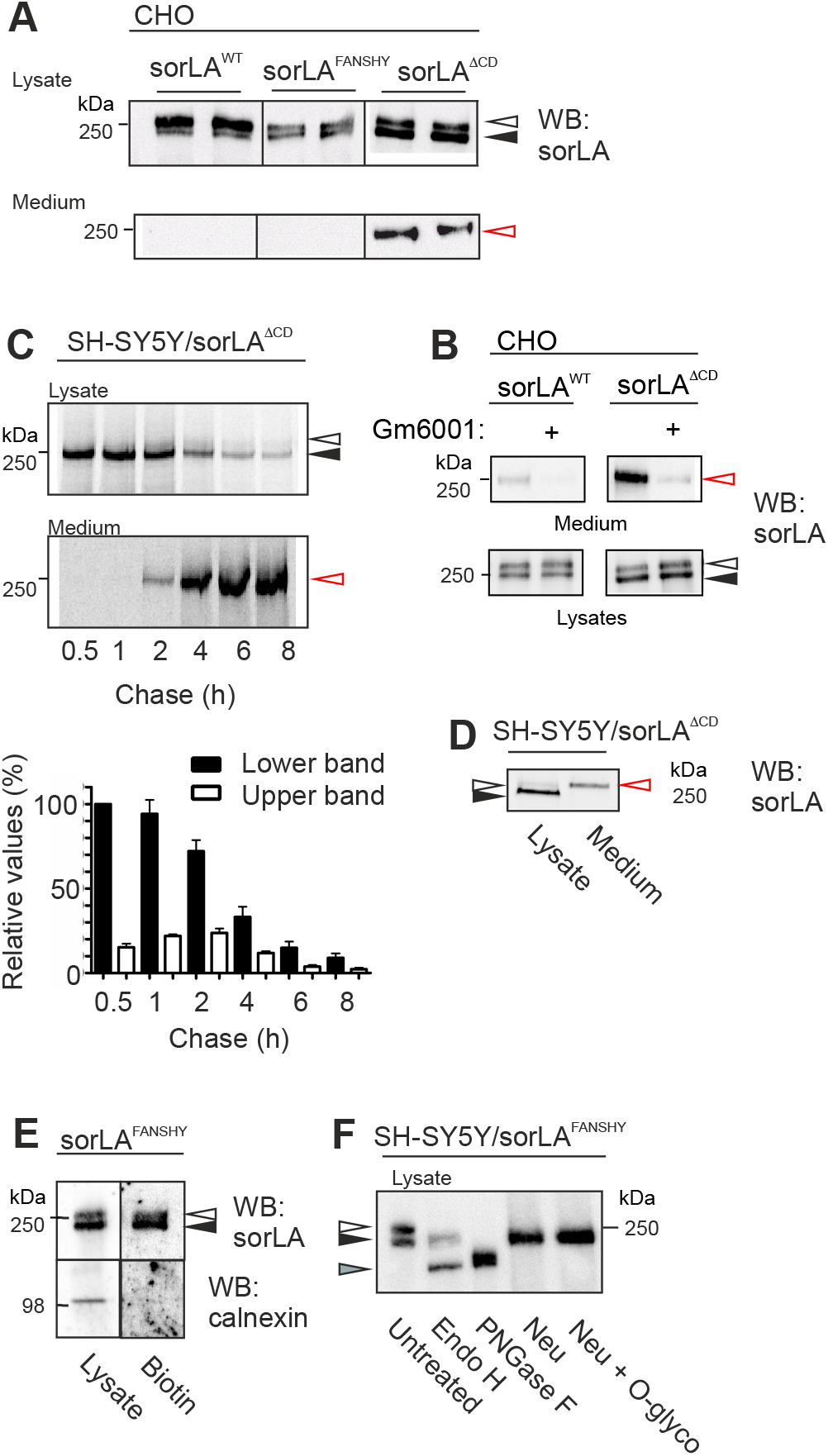
Correlation between complex-type *N*-glycans, shedding, and intracellular trafficking of sorLA mutants. **(A)** WB analysis of lysates and medium from transfected CHO cells that express sorLA^WT^, sorLA^FANSHY^ or sorLA^ΔCD^ using a polyclonal antibody against the luminal receptor domain. **(B)** Cells treated with the TACE inhibitor Gm6001 (+) shows strongly reduced shed sorLA in the medium, accompanied by accumulation of the upper variant in lysates, most prominently observed for cells expressing sorLA^ΔCD^ as these undergo much faster processing than sorLA^WT^. **(C)** SH-SY5Y cells expressing sorLA^ΔCD^ lacking the cytosolic tail were subjected to a pulse-chase protocol in order to follow maturation and shedding over an 8 h interval. Cells and medium were harvested at the indicated time points, and sorLA^ΔCD^ proteins were immunoprecipitated and subsequent analyzed by SDS-PAGE and radiofluorography. Quantification of mature (white) and immature (black) sorLA^ΔCD^ from lysates is shown from three independent experiments is shown below. **(D)** WB analysis of lysates and medium from SH-SY5Y cells expressing sorLA^ΔCD^ indicates loss of mature variant in the lysate caused by shedding of the high-molecular variant. **(E)** Cell surface proteins from SH-SY5Y cells expressing sorLA^FANSHY^ were labeled with biotin and precipitated using Streptavidin coated beads. The biotinylated proteins together with total lysates were analyzed using SDS-PAGE and WB for sorLA and calnexin. **(F)** Lysates from SH-SY5Y cells expressing sorLA^FANSHY^ were enzymatically digested with Endo H, PNGase F, and either neuraminidase (Neu) alone or in combination with *O*-glycosidase. The homogenates were subsequently separated by SDS-PAGE and analyzed by WB.

SorLA^FANSHY^ represents a variant with Ala substitutions of the ^2172^FANSHY motif located in the CD, which disrupts binding to the retromer subunit VPS26 with subsequent receptor accumulation in the tubular endosomal network (TEN) and less receptor undergoing retrograde trafficking (16, 46). This mutant is highly relevant for understanding the role of sorLA in Alzheimer’s disease, since the only naturally occurring variants of sorLA yet identified with disease-associated mutations in the cytoplasmic domain, all affect amino acids in this motif:, p.Ala2173Thr, p.Asn2174Ser, p.Ser2175Arg, and p.His2176Arg) (47–49). We previously showed that the SorLA^FANSHY^ mutant strongly reduces (>60%) ectodomain shedding in SH-SY5Y cells (16), and here we also showed that this mutant in CHO cells resulted in a very low level of shed ectodomain (below detection limit) (Fig. 6A). WB analysis of the cell lysates confirmed that sorLA^FANSHY^ underwent less *N*-glycan maturation with the two glycoform bands being present with similar intensity (Fig. 6A). Both bands were present at the cell surface (Fig. 6E), and deglycosylation analysis suggested an otherwise similar difference in mainly sialylated *N*-glycans between the upper and lower variant of sorLA^FANSHY^ (Fig. 6F). These data are in line with a model where sorLA has to undergo retrograde trafficking for acquiring maturation of *N*-glycans to complex-type.

### *In vivo* confirmation that sorLA maturation rely on retromer-associated retrograde/endosomal trafficking

Previous studies identified the VPS26 subunit of retromer as an interaction partner for sorLA (16), and the sorLA^FANSHY^ mutant impairs binding to the retromer subunit VPS26 required for retrograde transport (16, 46). In order to provide *in vivo* support for our hypothesis that sorLA maturation relies on retromer-dependent trafficking from the endosome, we studied sorLA maturation in a mouse model deficient in the retromer subunit VPS26. We tested sorLA expression by Western blot analysis in hippocampal brain homogenates from 7 month (Fig. 7A, B) and 13-14 month (data not shown) old mice, and observed a specific decrease in the levels of mature sorLA, whereas no change was observed for the immature receptor form. This shows that endogenous sorLA failed to undergo maturation when retrograde transport is blocked *in vivo* in mice, supporting our hypothesis that retromer dependent retrograde transport is essential for receptor maturation.

**Fig. 7.**
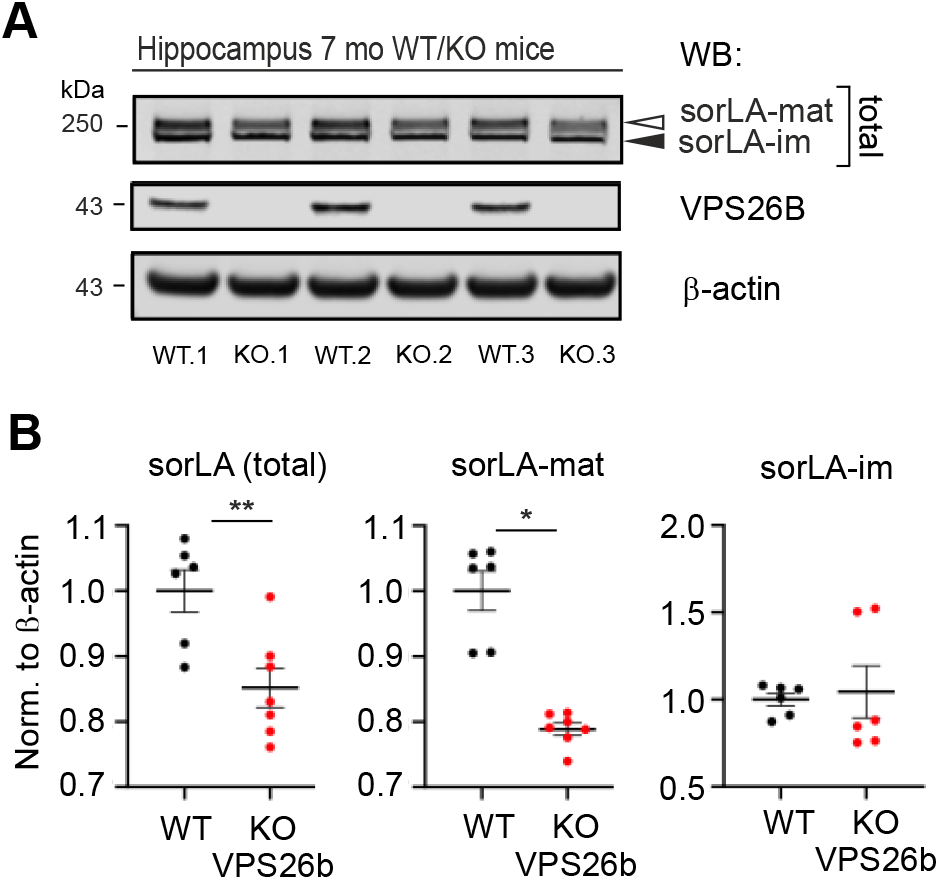
Endogenous sorLA fails to mature *in vivo* in brains of VPS26b knockout mice. **(A)** WB analysis of samples from 3 wildtype (WT) and 3 VPS26b knockout (KO) hippocampal homogenates probed with antibodies for sorLA, VPS26B, and β-actin. Signals for sorLA-mat and sorLA-im are indicated with white and black arrowheads, respectively. **(B)** Quantification of total receptor expression as well as the separate mature and immature forms of endogenous sorLA from WB analysis presented in panel A, showing a significant decrease (corresponding to 21%) in sorLA maturation in the VPS26b-deficient (KO) mice (N=7) compared to wildtype (WT) control mice (N=6). Data expressed as mean ± S.E.M, from two independent experiments.

## DISCUSSION

Our study provides an unusual model for the trafficking of sorLA in the secretory pathway schematically illustrated in Figure 8, where we propose that full maturation of sorLA with complex *N*-glycosylation occurs only following first transport to the cell surface and endocytosis, and that these steps are required for ectodomain shedding of sorLA. We propose that sorLA on its first pass to the cell surface can escape the typical Golgi *N*-glycan maturation to complex-type perhaps through interactions with cargo, which may relate to sorLA’s ability to chaperone APP through the secretory pathway and bypass amyloidogenic processing in the endosome (50). This may be in agreement with our previous finding that sorLA directs complexed APP into retrograde transport and affects the glycosylation of APP (19). We showed that retrograde trafficking of sorLA and acquisition of complex-type *N*-glycans was a requisite for TACE-mediated ectodomain shedding (Fig. 5), suggesting that sorLA before or during retrograde transport changes molecular state (or location) such that part of its *N*-glycans can be converted in a process initiated by MGAT1 to complex-type sialylated structures. Moreover, we demonstrated that the conversion of *N*-glycans is required for TACE-mediated shedding, suggesting that sorLA undergoes further molecular state changes by acquiring complex *N*-glycans (Fig. 3). Our proposed model can explain why shed sorLA exists as one glycoform in CSF and cell culture medium, while sorLA exists as two distinct glycoforms in homogenates of brain and cultured cells (7, 14–17).

**Fig. 8.**
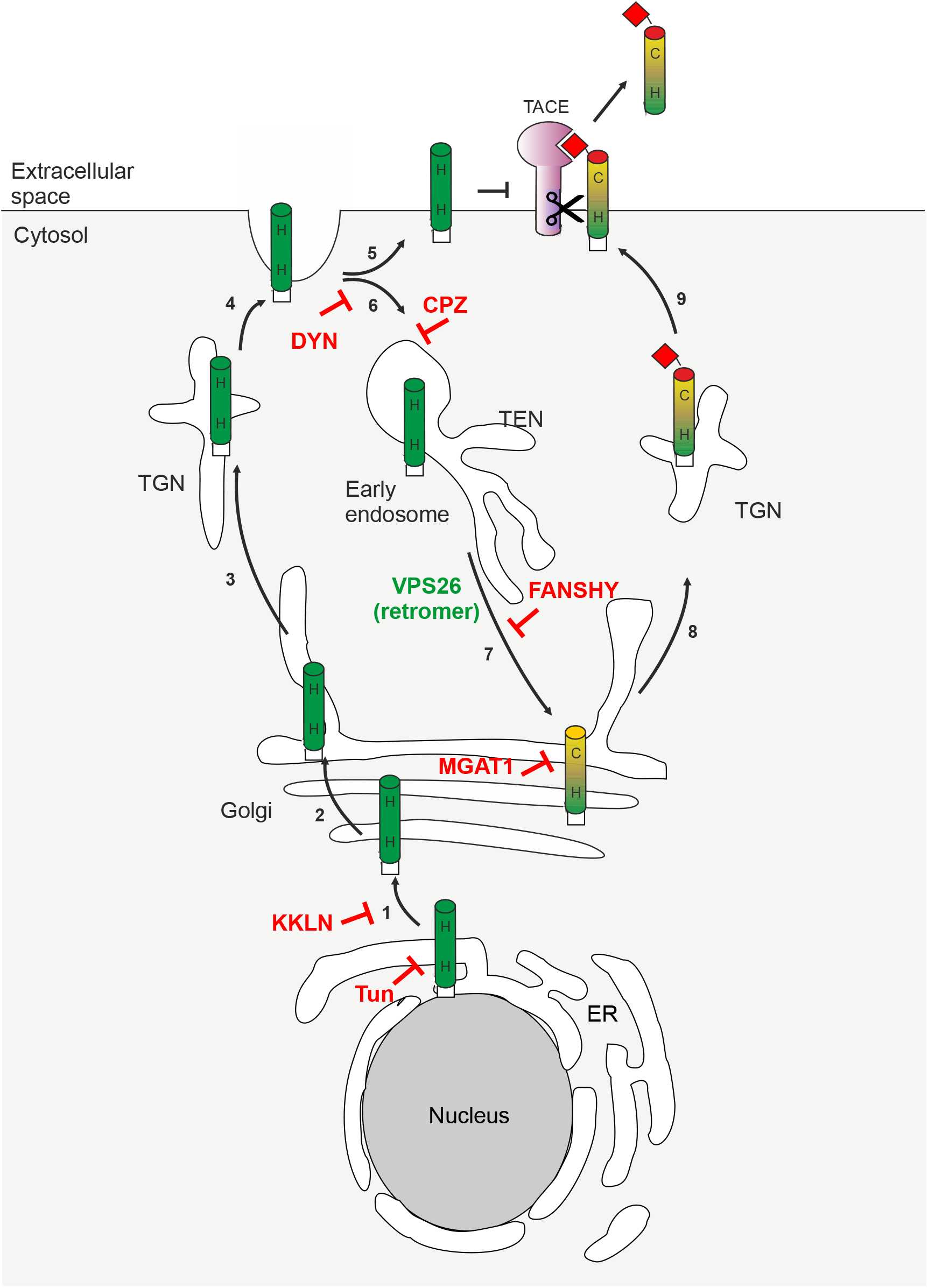
Proposed model for sorLA trafficking, *N*-glycosylation and shedding. Newly synthesized sorLA undergoes glycosylation in the ER and after transfer to the cis-Golgi (1), immature sorLA is able to escape maturation of high-mannose (H) *N*-glycans (2), and pass to the cell surface (3, 4). SorLA with the immature *N*-glycans is not substrate for TACE (5), but can internalize via clathrin-coated pits (6) to early endosomes from where it undergoes retrograde trafficking in a retromer-dependent route to the Golgi (7). Here *N*-glycans can mature to complex-type (C) that return to the cell surface (8, 9) and mature sorLA is then accessible to TACE cleavage and shedding.

SorLA is a large (2,214 amino acids) heavily *N*-glycosylated (28 potential *N*-glycosites) type-1 transmembrane glycoprotein (Fig. 2A, D). We clearly demonstrated that sorLA at the cell surface exists as two distinct *N*-glycoforms differing in sensitivity to endo H digestion and implying that only one of these have acquired complex-type structures (Fig. 2A, B). It is highly challenging to isolate sorLA and perform direct structural analysis of the many *N*-glycans, and even more so to obtain site-specific information of sorLA at the cell surface. We therefore took advantage of gene engineering (KO of *MGAT1)* to dissect the glycosylation process, and demonstrate that the first committed step for complex-type *N*-glycosylation directed by MGAT1 was essential for TACE-mediated shedding (Fig. 3D, E). We further used gene engineering (KO of *ST3GAL3/4/6)* to show that sialylation of *N*-glycans was not required. The predicted structural outcomes of the gene engineering were previously validated (24), and the results clearly demonstrate the value of genetic dissection strategies to probe functional importance of distinct glycosylation features in cell model systems.

The unusual coupling between *N*-glycan maturation and retrograde trafficking route proposed for sorLA is based on complementary results obtained with inhibitors, mutant constructs and a retromer-subunit knockout mouse model (Fig. 8). It is well established that sorLA internalizes via clathrin-coated pits and undergoes retrograde sorting in the endosome to Golgi in a process dependent on a cytosolic tail ^2172^FANSHY motif that binds to VPS26 of the retromer complex (16). We found that blocking retrograde transport by either DYN or CPZ blocked the maturation of *N*-glycans on sorLA (Fig. 4, 5), and this was recapitulated with sorLA mutants deficient in this cytosolic motif (Fig. 6). Furthermore, we established how retromer activity is a prerequisite for proper sorLA maturation, using hippocampal homogenates from VPS26b-deficient mice. While glycosylation of glycoproteins during recycling or retrograde transport has been reported, previous studies have mainly found this limited to resialylation of glycans that could be attributed to completion of glycosylation or in response to neuraminidase desialylation at the cell membrane (51). To our knowledge conversion of high-mannose to complex-type *N*-glycans during retrograde transport has not been reported for other glycoproteins, and it has been debated if retrograde trafficking extends all the way to the early cis-Golgi where MGAT1 directs this fundamental transformation of *N*-glycans (52).

Our results confirm previous studies implicating TACE as the metallo-proteinase responsible for shedding of the sorLA ectodomain (14, 26, 27). This process is regulated by receptor-interactions of sorLA with selected ligands, including head-activator as well as the tetraspanin CD9 that increase sorLA shedding (14, 53). Phosphorylation of the cytosolic domain of sorLA also increases sorLA shedding (15). Our study now adds another regulatory mechanism for sorLA shedding being acquisition of complex-type *N*-glycans. Interestingly, we found that action of MGAT1 was essential while the capacity for sialylation of *N*-glycans was not. How the state of the *N*-glycans on sorLA affects TACE processing is currently unclear, but *N*-glycans on sorLA are found close to the juxtamembrane region where TACE cleaves (Fig. 2D), and glycans including *O*-glycans in these regions are well known to be able to co-regulate ectodomain shedding (54). It is also conceivable that the change in glycosylation state affects molecular interactions with other proteins and enable TACE action.

SorLA is strongly implicated in AD pathogenesis and APP processing (22, 55, 56). Interestingly, gene variants that lead to substitution of amino acids in the ^2172^FANSHY motif are so far only identified in AD patients and not in non-AD controls (47, 48). This suggests that retromer-dependent trafficking is essential for sorLA to perform its protective role in development of AD in agreement with retromer dysfunction leading to core pathological features of AD (57), and defective endosomal trafficking suggested as a culprit of AD (58). Recently, a family with high incidence of AD carrying a variant locus at a *N*-glycosylation site in the sorLA VPS10p domain was described, adding further interest to the role of glycosylation in regulating functions of sorLA (59).

In contrast to a range of risk-factor genes, *SORL1*-- together with *APP, PSEN1* and *PSEN2*-- is now generally acknowledged as only the fourth gene that is causally pathogenic in AD. Gene association studies have identified many variants in *SORL1*, and ongoing studies are expected to increase this list, and in the absence of functional insight into the large and complex sorLA glycoprotein it is challenging to identify and assess deleterious variants and their roles in disease. The present results provide a detailed map of the trafficking itinerary of sorLA and its glycosylation and maturation processes that will aide in mechanistic insight into sorLA variants and their linkage to AD. Furthermore, our study points to novel approaches for drug discovery through screening for sorLA maturation as well as biomarker discovery with shed sorLA in CSF and other bodyfluids.

## Supporting information

Suppmental Figures

## ACKNOWLEDGEMENTS

Marit Nyholm Nielsen and Sandra Bonnesen are thanked for excellent technical assistance. Funding for this work was kindly supported by the Novo Nordisk Foundation, the Augustinus Foundation, the Hørslev Foundation, the Hartman Foundation, the Henrik Henriksen Foundation, the Lundbeck Foundation, and the Danish National Research Foundation (DNRF107). We are also grateful to Drs. Sang-Rae Lee and Young-Hyun Kim for sharing the VPS26b KO mice.

## AUTHOR CONTRIBUTION

SC, YN, SS, CG, CV, SAS, HC, and OA all contributed to design and analyze the conducted experiments. HC, SC, SAS, and OA wrote the manuscript.

