## Supplementary material for "Endosomal trafficking is required for glycosylation and normal maturation of the Alzheimer’s-associated protein sorLA": Suppmental Figures

SUPPORTING INFORMATION

**Fig. 1S. Immature sorLA migrates indistinguishable from sorLA$^{KKLN}$**

WB analysis of sorLA$^{WT}$ or sorLA$^{KKLN}$ using either 2 or 20 μg of lysates to document the presence of immature receptor by addition of the KKLN ER-retention motif.

**Fig. 2S. Identification of immature and complex sorLA glycosylations using lectin blotting.**

SorLA proteins were immunoprecipitated (IP) from lysates of SH-SY5Y cells expressing sorLA$^{WT}$, sorLA$^{FANSHY}$, sorLA$^{\Delta CD}$ (left) or soluble sorLA$^{WT}$ from medium (right) with an antibody against the extracellular receptor fragment. The precipitated proteins were separated by SDS-PAGE and analyzed by Western blotting (WB) for sorLA or by Lectin blotting (LB) for the presence of specific glycans. The lectins Concanavalin A (ConA), Wheat Germ Agglutinin (WGA), Ricinus Communis Agglutinin 120 (RCA120), Sambucus Nigra Lectin (SNA).

**Fig. 3S. Amino acid sequence of included sorLA tail variants**

**Fig. S4. Endogenous sorLA fails to mature *in vivo* in brains of VPS26b knockout mice.**

**(A)** Representative WB analysis of samples from 2 wildtype (WT) and 2 VPS26b knockout (KO) hippocampal homogenates of mice 13/14 mo old probed with antibodies for sorLA, VPS26B, and β-actin. Signals for sorLA-mat and sorLA-im are indicated with white and black arrowheads, respectively.

**(B)** Quantification of total receptor expression as well as the separate mature and immature forms of endogenous sorLA from WB analysis presented in panel A, showing a significant decrease (corresponding to 24%) in sorLA maturation in the VPS26b-deficient (KO) mice (N=4) compared to wildtype (WT) control mice (N=5). Data expressed as mean ± S.E.M.

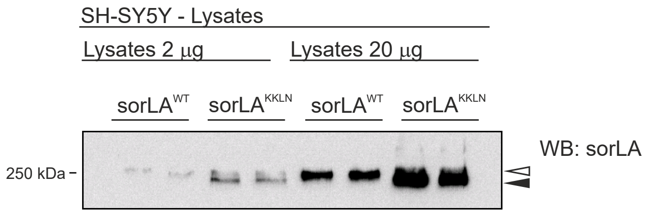


*Supporting Information*, Fig. S1

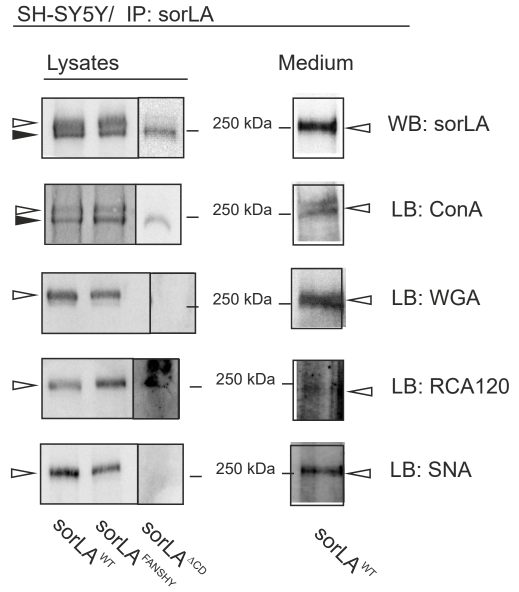


*Supporting Information*, Fig. S2

| | |
|---|---|
| SorLA$^{WT}$ | KHRRLQSSFTA**FANSHY**SSRLGSAIFSSG**DDLGEDDED**APMITGFSDDVPMVIA |
| ΔCD | KHRRLQSS |
| FANSHY | KHRRLQSSFTA**A**A**AAAA**SSRLGSAIFSSGDDLGEDDEDAPMITGFSDDVPMVIA |
| KKLN | KHRRLQSSFTAFANSHFSSRLGSAIFSSGDDLGEDDEDAPMITGFSDDVPMVIA**KKLN** |

*Supporting Information,* Fig. 3S

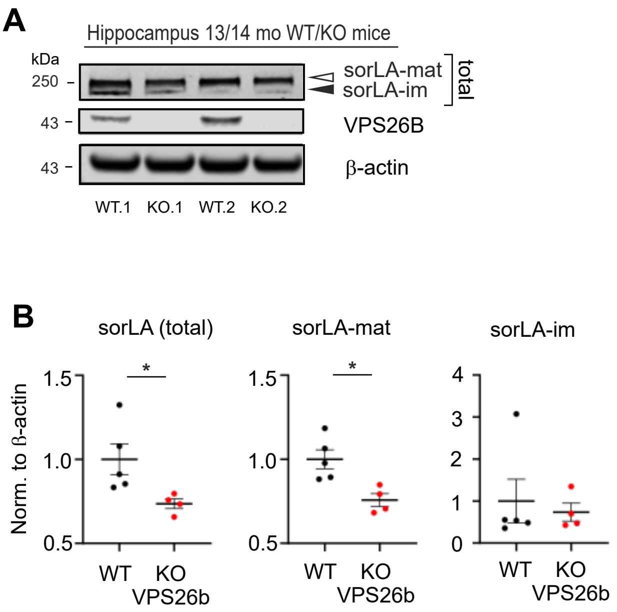


*Supporting Information*, Fig. S4
